# LncPTPred: Predicting lncRNA-Protein Interaction based on Crosslinking and Immunoprecipitation (CLIP-Seq) Data

**DOI:** 10.1101/2025.01.07.631626

**Authors:** Gourab Das, Troyee Das, Zhumur Ghosh

## Abstract

Long non-coding RNA (lncRNA)-Protein Interaction (LPI) across diverse biological systems directly and indirectly regulates various cellular processes. Experimental assays to recognize the protein binding partners of lncRNAs is highly time consuming and expensive. *In-silico* predictive approaches involving pattern recognition technique provides a promising alternative to it by reducing the search space. Our work identifies such hidden pattern from within the Cross-linking immunoprecipitation sequencing (CLIP-Seq) data which aids to overcome the problem of obtaining real negative dataset and thus offers a state-of-the-art machine learning (ML) based prediction algorithm to predict LPI. Initial phase of this work involves preparation of the training dataset and the next phase is devoted towards developing the ML based model to perform prediction operation. In order to show the efficacy of our model, its performance has been compared against that of the contemporary prediction tools, where the result clearly shows the outperformance of our model. Moreover it also provides the segments of interaction within the lncRNA loci which acts as a roadmap for precise designing of the validation experiment. The LncPTPred tool has been provided in terms of web server as well as standalone version in github.

**Web server Link:** http://bicresources.jcbose.ac.in/zhumur/lncptpred/

**Github Link:** https://github.com/zglabDIB/lncptpred.git

## INTRODUCTION

Long noncoding RNAs (lncRNAs) play vital role towards regulating a wide spectrum of biological processes including gene transcription, epigenetic modifications along with other post-transcriptional regulatory mechanisms. These versatile molecules are poorly conserved and adopt various mechanisms to implement such regulatory actions which includes RNA-RNA(1), RNA-protein (2) and competing endogenous RNA (ceRNA) (3) mode of interaction. Among these, lncRNA-protein interaction(LPI) constitutes one such important mode of interaction which not only have implication in various normal biological processes and different disease systems but also influence lncRNA processing, their modifications, stability and function (4). The advancement of next gen sequencing techniques have facilitated the detection of RNA-protein interaction viz. cross-linking immunoprecipitation-seq (CLIP-seq) (5) and RNA immunoprecipitation seq (RIP-seq) (6), RIP-Chip etc. However, irrespective of the nature of the experiments, they are costly and time-consuming. Alternatively, *in-silico* approaches like Molecular docking(MD) and ML based approach have been adopted for quick screening of potential RNA-protein interaction, more specifically LPIs. But MD being computationally expensive have necessitated the introduction of Artificial Intelligence (AI) based computational tool to predict the LPI binding interactions. Thus Machine learning (ML) or Deep learning(DL) based approaches have gained importance which provides a time optimized solution.

There have been reports on ML based RPI prediction models incorporating either sequence based or structure based features. Initial study by Pancaldi et al. (7) mentioned mRNA-protein target prediction using Random Forest (RF) and Support Vector Machine (SVM) models. They have incorporated various features associated with the genes and proteins of the *Saccharomyces cerevisiae* (yeast). Muppirala et al. (8) created RPISeq using same RF and SVM models with 3-mer protein and 4-mer RNA features from the sequences. Wang et al. (9) trained Naïve-Bayes classifier for RPI and Lu et al. (10) proposed lncPro to generate association score using Fisher linear discriminant analysis (11). RPI-Pred (12) model made use of both sequence and structural features to train SVM classifiers whereas IPMiner (13) used deep learning based stacked autoencoder to create computational RPIs. Some of the recent studies related to LPI prediction includes Human lncRNA-protein interactions (HLPI)-Ensemble (14) based ensemble models in terms of RF, Extreme Gradient Boosting (XGBoost) and SVM based general model. Li et al. (15) proposed Capsule-LPI tool incorporating multimodal features in terms of secondary structure, motif, sequence and physiochemical properties for both lncRNA and protein where Capsule network forms the basis of their prediction model. LPI-CNNCP (16) used Convolution Neural Network (CNN) model to perform prediction task where the feature set was generated using copy padding trick to convert variable length sequences of lncRNA/protein into fixed length inputs, incorporating one-hot encoding in order to transform CNN-compatible image data. LPI-HyADBS (17) formed hybrid network containing deep neural network (DNN), XGBoost, and SVM with a penalty Coefficient of misclassification (C-SVM) to execute prediction strategy by incorporating Pyfeat (18) based biological features, extracted from lncRNA and proteins sequence. EnANNDeep (19) constructs ensemble framework containing adaptive k-nearest neighbor classifier, deep neural network, and deep forest to perform prediction task where features from multiple sources are integrated to determine lncRNA-protein pair. The most predominant datasets used as training data by these specified models are NPInter (20), PRIDB (21) & RPIs (8,13) which contain only real positive interaction pairs.

It is important to emphasize that the biggest pitfalls corresponding to most of these models is the non-availability of real non-LPI dataset or real negative dataset. Negative dataset were generated by random shuffling of unlabelled interactions. Thus, it induces noise within trained models that leads to inaccurate learning. In this work, we have developed LncPTPred; an LPI prediction tool to address this issue by generating properly labeled dataset yielded from CLIP-Seq based biological assays consisting of both Photoactivatable Ribonucleoside-Enhanced Crosslinking and Immunoprecipitation (**PAR-CLIP**) (22) and High-throughput sequencing of RNA isolated by crosslinking immunoprecipitation (**HITS-CLIP**) (23). Guiding principle corresponding to our training data generation procedure considers that RNA binding Protein found from PAR-CLIP and HITS-CLIP sequences are positive data and rest of the non-interacting segments are the negative one. Moreover, it is able to predict the exact location of interaction which is not provided by any other state-of-the-art prediction tools mentioned above. Further, the performance of LncPTPred has been compared against other state-of-the-art LPI prediction models. Finally, we have tested our models deployed for LncPTPred by correctly predicting well known LPIs in case of certain diseases. User can access LncPTPred from http://bicresources.jcbose.ac.in/zhumur/lncptpred/. It’s standalone version is available at https://github.com/zglabDIB/lncptpred.git.

## MATERIAL AND METHODS

The detailed workflow associated with LncPTPred depicted in **Figure 1** can be segregated into two main modules. **Initial module deals** with **Data Curation** where LPI data has been extracted from PAR-CLIP and HITS-CLIP based CLIP-Seq assay. Numerous preprocessing operations have been incorporated in order to filter out lncRNA sequence segments from the list of all binding sequences. **The second module** is dedicated towards developing Feature Creations and **Machine Learning Models** where two of the main sub-modules include development of stacking classifier based Meta learner using **scikit-learn** (https://scikit-learn.org/stable/) (24) and **Hyperparameter Optimization** using **Optuna** (25). Finally there is the Output Module and data visualization part which provides the details regarding the interacting lncRNA segment. Information on each module as well as curated datasets is detailed below.

**Figure 1:**
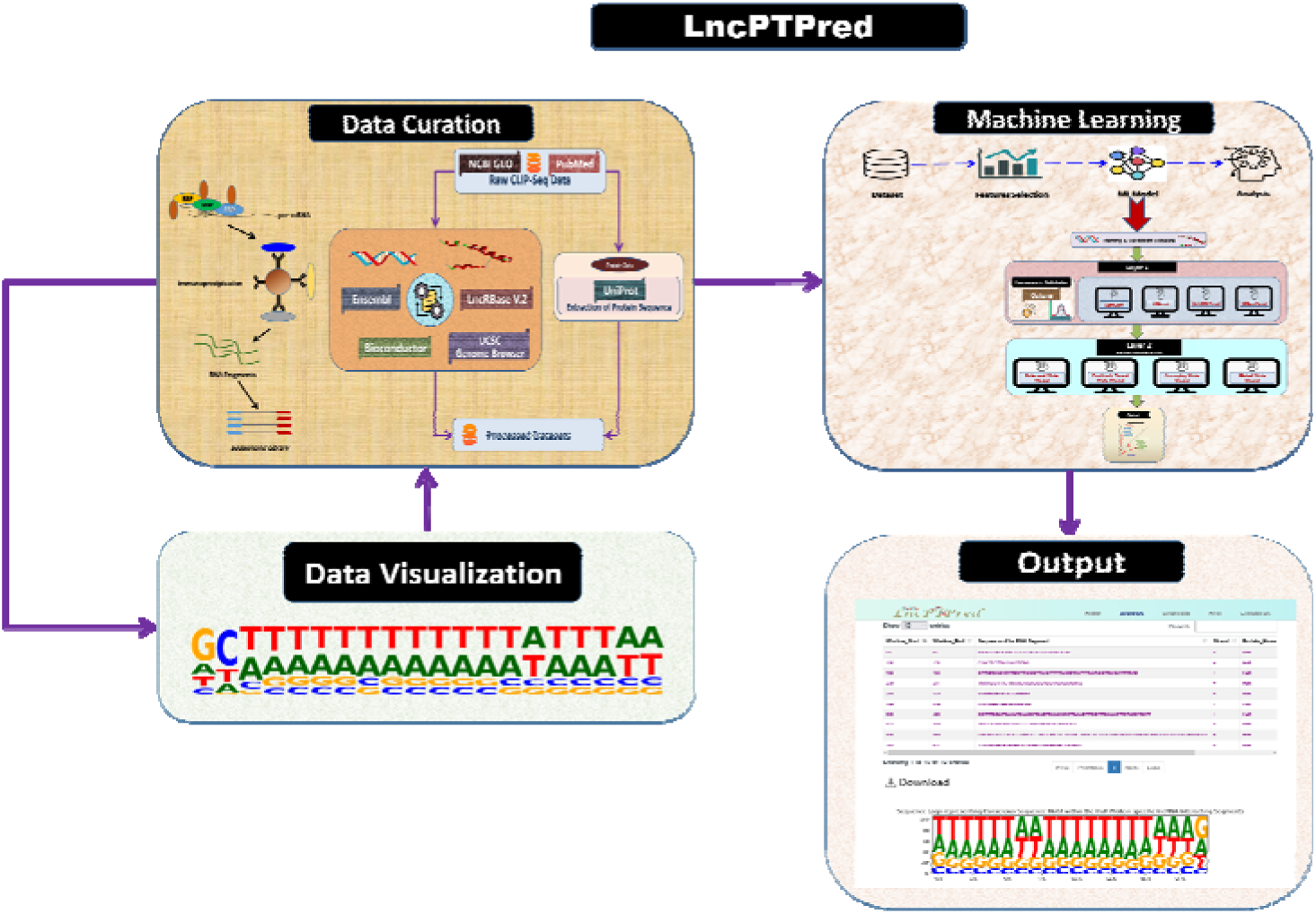
Workflow of the entire work. The flow diagram depicts the various steps involved towards development of LncPTPred which includes Data Curation from CLIP-Seq data, development of the ML based models, output generation and data visualization

### Dataset

Given the maximal importance towards the generation of properly labelled dataset we have considered CLIP-Seq experimental datasets viz. PAR-CLIP (22) and HITS-CLIP (23) data. For a particular protein, protein bound RNA fragments have been considered as positive data and rest of the non-bounded fragments have been considered as real negative data. These data have been curated from GEO (https://www.ncbi.nlm.nih.gov/geo/) and pubmed ID: 22081015, as provided in **Supplementary File** 1 in detail. The entire set of GSE numbers, the pubmed ID of the input datasets constituting the training and validation data along with the protein names and cell line information is provided in **Table 1**.

**Table 1:** Input Training Dataset utilized for developing LncPTPred.

| GSE_NUMBER/Pubmed_Id | Protein_Name | Cell_Line |
| --- | --- | --- |
| GSE38201 | ALKBH5 | HEK293 |
| GSE98856 | G3BP1 | HeLa |
| GSE160948 | SF3B1 | MDA-231-LM2 |
| GSE46908 | LIN28B | HEK293 |
| GSE38201 | C17orf85 | HEK293 |
| GSE102320 | HuR | 293T |
| GSE111900 | SRSF2 | Human erythroid leukemia (HEL) cell line |
| GSE130867 | SRSF1 | H1299 |
| GSE133724 | FMR1 | HEK293 |
| GSE160949 | SNRPA1 | MDA-231-LM2 |
| GSE98836 | EWSR1 | HeLa |
| GSE173628 | ESR1 | MCF7 |
| GSE38201 | CAPRIN1 | HEK293 |
| GSE38201 | C22orf28 | HEK293 |
| GSE62115 | MSI2 | K562 |
| GSE44615 | LIN28A | HEK293 |
| GSE44615 | LIN28B | HEK293 |
| GSE47078 | RC3H1 | HEK293 |
| GSE58352 | YTHDC1 | HeLa |
| GSE111188 | QKI | Mesenchymal-like HMLE cells |
| GSE39872 | LIN28A | H9 |
| GSE38201 | ZC3H7B | HEK293 |
| GSE63591 | YTHDF1 | HeLa |
| GSE49339 | YTHDF2 | HeLa |
| GSE19323 | PTBP2 | HeLa |
| GSE52977 | AUF1 | HEK293 |
| GSE46705 | METTL14 | HeLa |
| GSE29943 | HuR | HeLa |
| GSE46705 | METTL3 | HeLa |
| GSE46705 | WTAP | HeLa |
| GSE150925 | YBX1 | HEK293 |
| GSE102336 | ZCCHC4 | Hela |
| GSE53184 | ZFP36 | HEK293 |
| Pubmed:22081015 | TAF15 | HEK293 |
| Pubmed:22081015 | FUS | HEK293 |

The unknown test data have been provided in **Table 2**. Both the dataset in Table 1 and Table 2 undergoes several pre-processing steps as done under the 1st Module named Data curation.

**Table 2:** Unknown Test Dataset utilized to validate the performance of LncPTPred against external models.

| GSE_NUMBER/Pubmed_Id | Protein_Name | Cell_Line |
| --- | --- | --- |
| GSE98856 | G3BP2 | HeLa |
| Pubmed:22081015 | EWSR1 | HEK293 |

### Data Curation

Datasets of various CLIP-Seq experiments mined from GEO database were in diverse formats such as excel, text, bed, bedgraph and bigwig. Moreover, different build version of human sequences like **hg38, hg19** or even **hg18** have been considered while generating the CLIP-seq data as detailed in **Supplementary File 1**. It necessitates the incorporation of robust pre-processing techniques in order to extract the protein bound RNA segments corresponding to a particular CLIP experiment. Furthermore, it is essential to screen out lncRNAs from the given set of protein bound fragments. The flowchart in **Figure 2** depicts the different steps(as detailed below) to generate properly labeled data to be utilized by ML based target prediction model.

**Figure 2:**
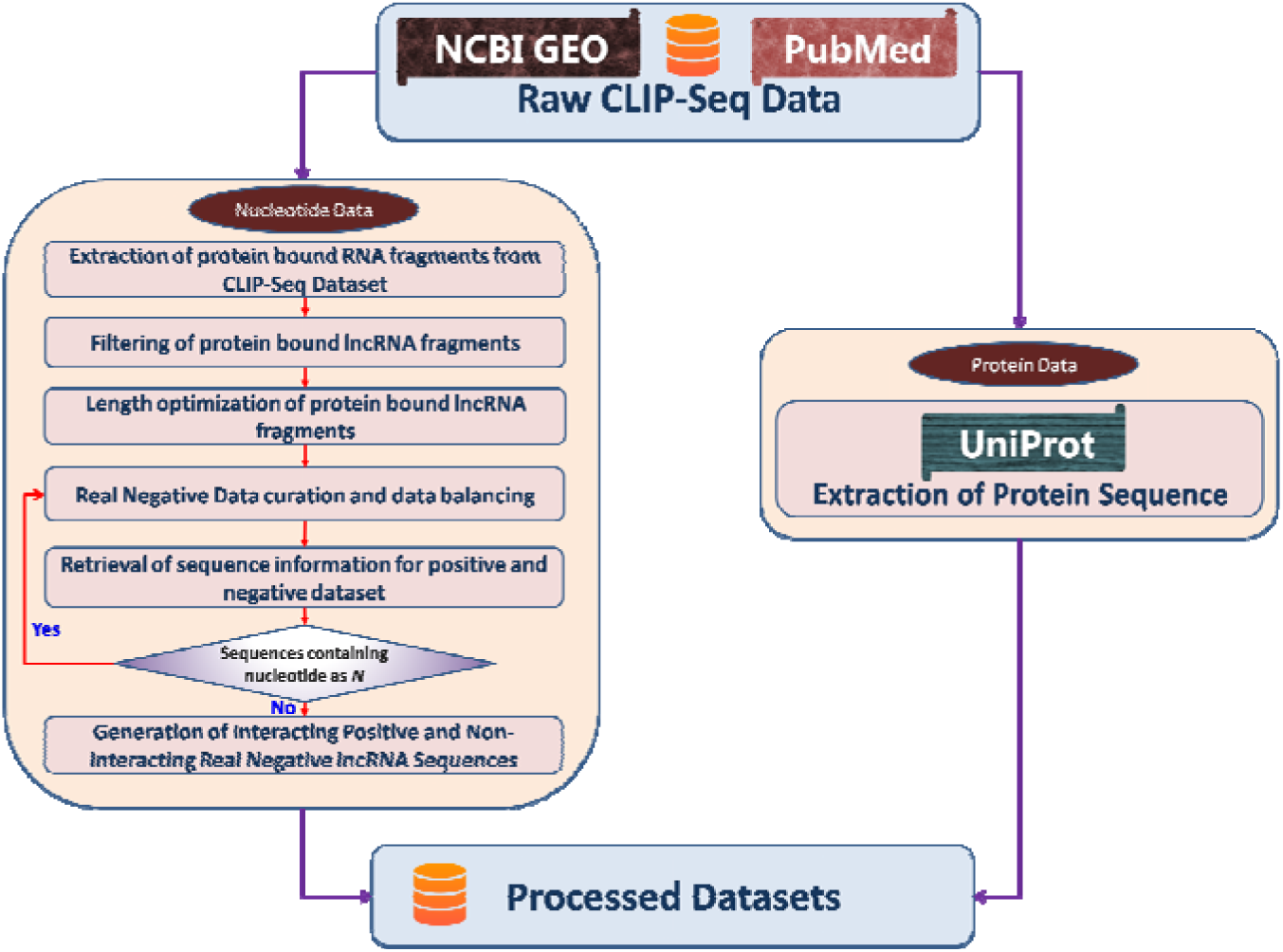
Data Curation Workflow. It depicts the pre-processing of the raw CLIP-seq data to build the training dataset, validation and unknown test dataset which includes the extraction of positive and real negative data. It also depicts the curation of protein sequence information

#### (a) Extraction of protein bound RNA fragments from CLIP-Seq Dataset

Protein bound RNA fragments have been extracted from the downloaded files in various formats viz. bed, bedgraph, excel, text and bigwig. More specifically the start & end location, strand and chromosome specific information of these protein bound RNA fragments have been obtained.

#### (b) Filtering of protein bound lncRNA fragments

The reference files were created at first, prior to mapping the protein bound RNA fragments with them. This has been done using Ensembl Biomart (26) and LncRBase V.2 (27) as the reference database corresponding to human genome version **hg38**.

Since some of the input dataset have been generated using genome build hg19 and hg18, UCSC genome liftover tool (http://genome-euro.ucsc.edu/cgi-bin/hgLiftOver) have been used to convert **hg38** reference file into **hg19 and** CrossMap (28) tool (https://genviz.org/module-01-intro/0001/06/02/liftoverTools/) to convert hg19 reference file to hg18. The reference files are further filtered with length cut off of <200 bases as lncRNA length is typically ≥ 200 bases. All the reference files contain chromosome number, strand and start & end position of the lncRNA loci.

Subsequently, the protein bound RNA fragments have been mapped to these reference files to filter out the protein bound lncRNA fragments. The pseudocode for performing the mapping at this step is as follows:

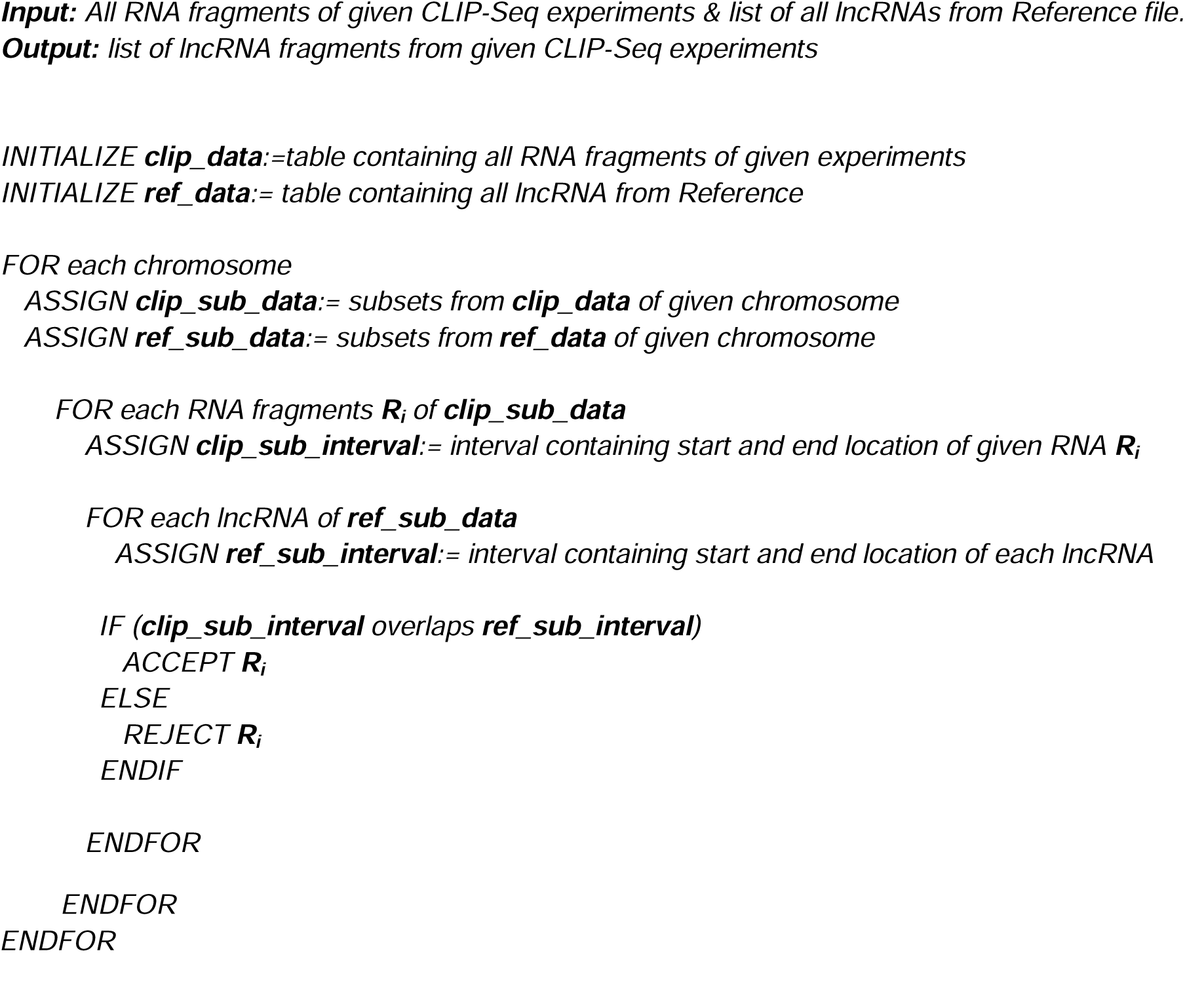

#### (c) Length optimization of protein bound lncRNA fragments

We next checked the distribution of the lengths of these protein bound lncRNA fragments in training, validation and unknown test dataset. Violin plot provided in **Figure 3** shows that the length of the protein bound lncRNA fragment size dominating around a threshold of ≤ 40 bases which is well supported by earlier reports (29). This is an important parameter for determining the window size of the search space to be selected by the users while using LncPTPred. In order to remove length bias without ignoring too much data segments we have filtered the length of lncRNA between 10 and 200 nucleotides.

**Figure 3:**
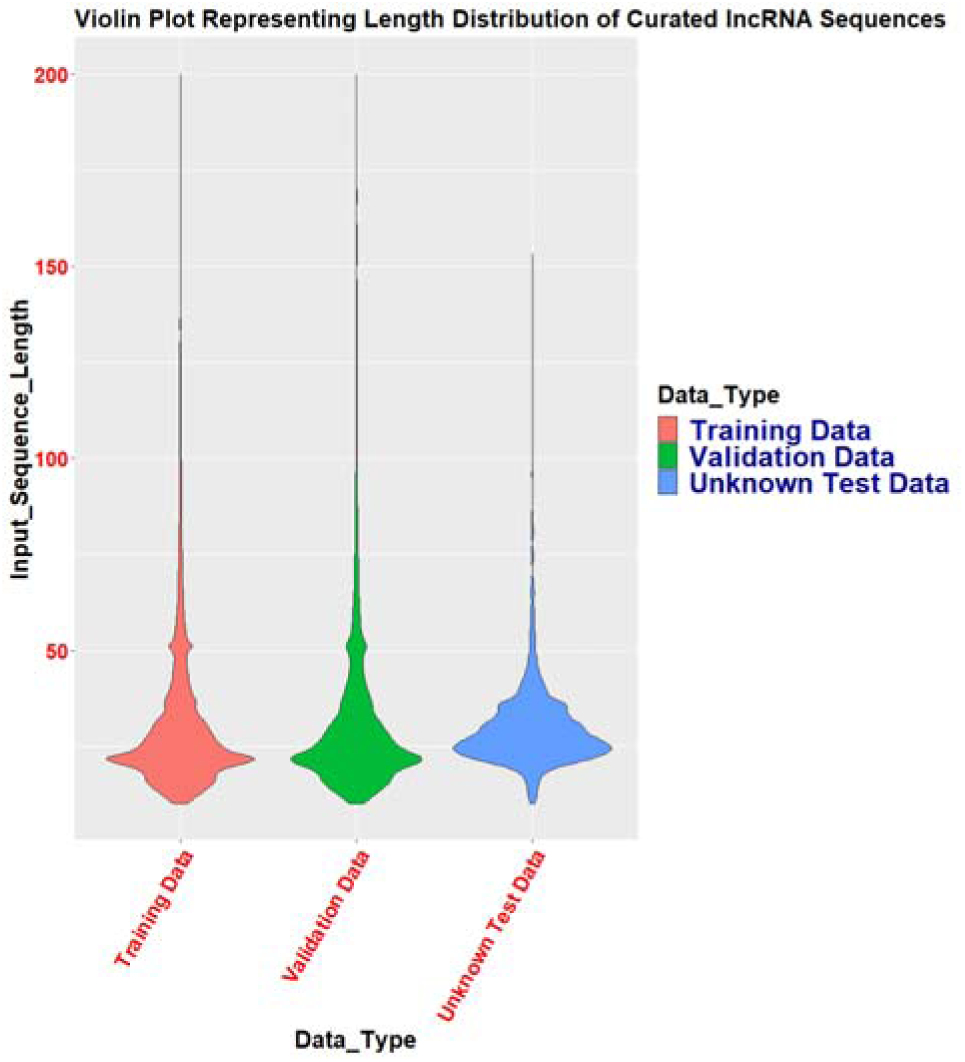
Length Distribution of Interacting lncRNA sequences extracted from CLIP-seq data. The violin plot shows the length distribution of interacting lncRNA sequences extracted from CLIP-seq data subdivided as training, validation and unknown test dataset which will serve as an important parameter for determining the window size of the search space to be selected by the users while using LncPTPred.

#### (d) Real Negative Data curation and data balancing

In a CLIP-Seq experiment for a particular protein, RNA fragments not bound to the protein have been considered as real negative data. In order to address data imbalance due to more number of negative data as compared to positive data, random sampling of non-bound lncRNA fragments (for a particular protein) have been done so as to maintain consistency with the positive dataset in terms of number, length and strands of the fragmented sequences. Corresponding to each positive dataset, we have generated negative data belonging to same chromosome and strands across 10KB to 1 MB cis-location on either side of the given positive data. The detailed pseudocode for negative data curation is given below:

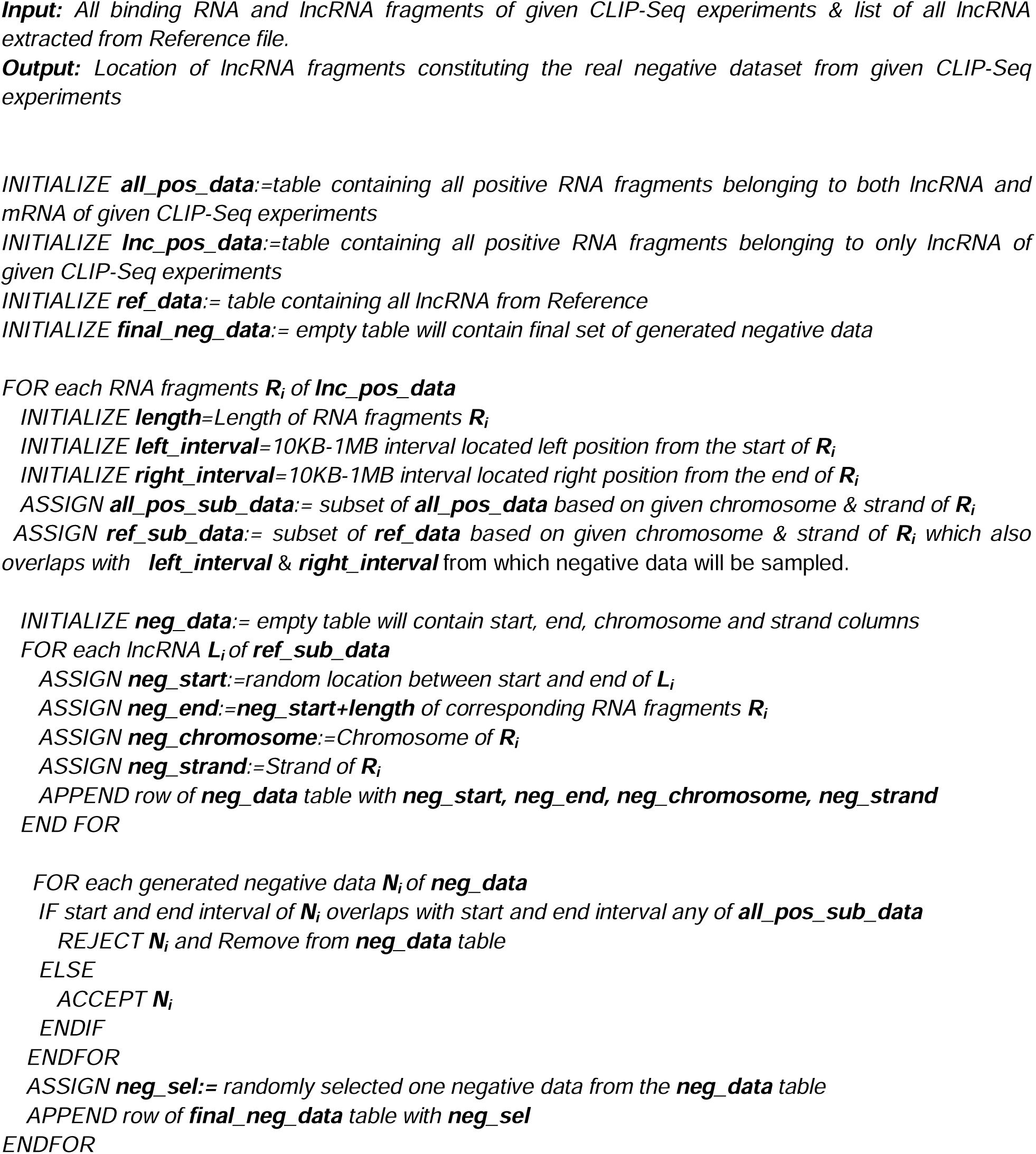

#### (e) Retrieval of sequence information for positive and negative dataset

We have extracted sequence information using R programming based **BSgenome** tool downloaded from **Bioconductor**(**30**) package manager. **BSgenome.Hsapiens.UCSC.hg38, BSgenome.Hsapiens.UCSC.hg19** and **BSgenome.Hsapiens.UCSC.hg18** packages have been deployed to extract the sequences corresponding to the hg38, hg19 and hg18 builds respectively based on given chromosome number, strand information, start & end location. Corresponding to the negative dataset, nucleotide bases for very few fragments remained missing and yielded **N (**for hg19 and hg18 build), which would have an adverse effect on the training procedures for ML operations further. Hence while retrieving the sequence information, presence of **N,** within the sequences for the negative dataset, necessitated repeating step **(d)** until all the fragments constituting the negative dataset are devoid of missing nucleotide **N.**

The final dataset to be used for ML training is provided in **Table 3**.

**Table 3:** Data Dimension corresponding to Training, Validation and Unknown Test Data.

| Dataset Type | # of Data |
| --- | --- |
| Training Data | 245501 |
| Validation Data | 27279 |
| Unknown Test Data | 9468 |

#### (f) Extraction of Protein Sequences

The sequences of 33 proteins corresponding to different CLIP-Seq experiments have been extracted from the Uniprot database (https://www.uniprot.org/)

### Machine Learning

This module consists of 3 subparts: **Data Preprocessing:** which generates ML compatible dataset**; ML Model Creation & Validation:** build stacking classifiers and train them with the ML compatible datasets; **Hyper Parameter Optimization (HPO):** tune numerous hyper-parameters associated with ML models using **Optuna** tool in order to enhance model performance

#### Data Preprocessing

This consists of two steps viz. Feature Selection & Data Normalization in order to extract the final set of features

### Feature Selection

Corresponding to both lncRNA and Protein sequences, features are created based on normalized count of nucleotide bases or amino acids described as follows:

### lncRNA Feature Selection

Selected features for lncRNA data are fundamentally embedded in their sequences containing strand specific information, normalized counts of unique nucleotides A, C, G, T; 2 contiguous nucleotides or di-nucleotides (AA, AC, AG, AT, …, etc.); 3 contiguous nucleotides (AAA, AAC, AAG, AAT, …, etc.) and 4 contiguous nucleotides (AAAA, AAAC, AAAG, AAAT, …, etc.); overall yielding 341 features as shown in **Table 4**.

**Table 4:**
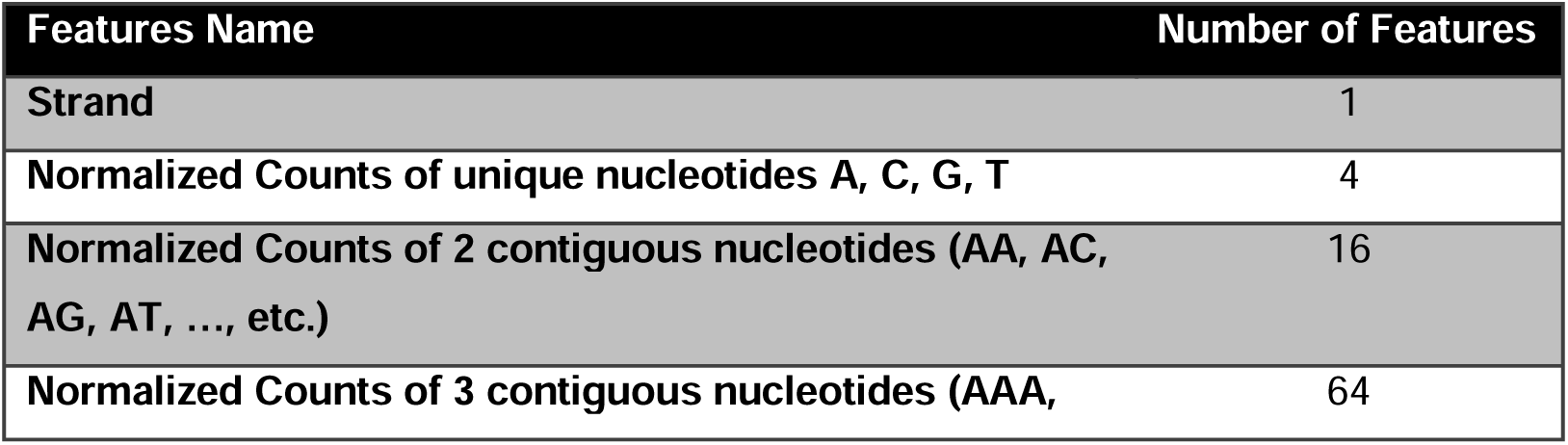
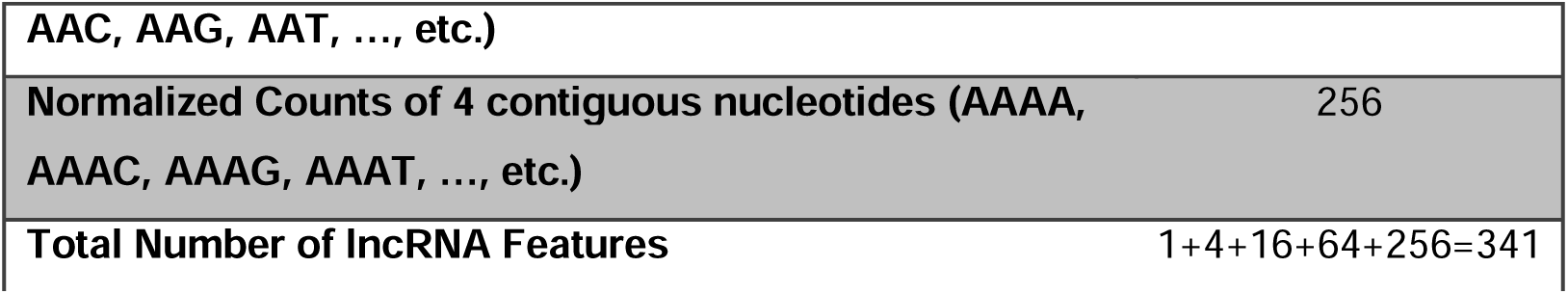
Detailed Feature Description extracted from raw lncRNA sequences from CLIP-seq dataset.

### Protein Feature Selection

Properties of amino acids are essential for generating the features from raw protein sequences. Here 20 primary amino acids and their important properties have been viz. **Hydropathy, Chemical, Charge and Polarity** along with **Volume** to be implemented in feature selection strategy. Most of the features (https://www.imgt.org/IMGTeducation/Aide-memoire/_UK/aminoacids/IMGTclasses.html) have been generated as specified by Pommié, C., et al (31). Amino acids can be classified into 3 types: **Hydrophobic (A, C, I, L, M, F, W, V), Neutral (G, H, P, S, T, Y) and Hydrophilic (R, N, D, Q, E, K)** corresponding to **Hydropathy** property based on Kyte-Doolitle scale (32). Using the normalized counts of single, di and tri **Hydropathy** grouping based strategies; total 39 features have been generated. Corresponding to the **Chemical, Charge and Polarity** based properties, amino acids are segregated into **Aliphatic (G, A, V, L, I, P), Aromatic (F, Y, W), Polar_Uncharged (S, T, N, Q), Sulpher_Contained (C, M), Positive_Negative_Charged (D, E, H, K, R).** Following the same procedure i.e. single, di and tri normalized counts, total 155 features have been generated. Finally, inclusion of **Volume** properties help us to classify the amino acids into 5 classes: **Very_Small (A, G, S), Small (N, D, C, P, T), Medium (Q, E, H, V), Large (R, I, L, K, M), Very_Large (F, W, Y)** ultimately resulting in generation of 155 more features following their normalized counts of occurrence. Besides, the normalized counts of individual amino acids additionally yields 20 more features resulting in an overall 369 feature. Accumulating set of all attributes from lncRNA and protein creates 710 features to be trained by ML models which have been detailed in **Table 5**.

**Table 5:**
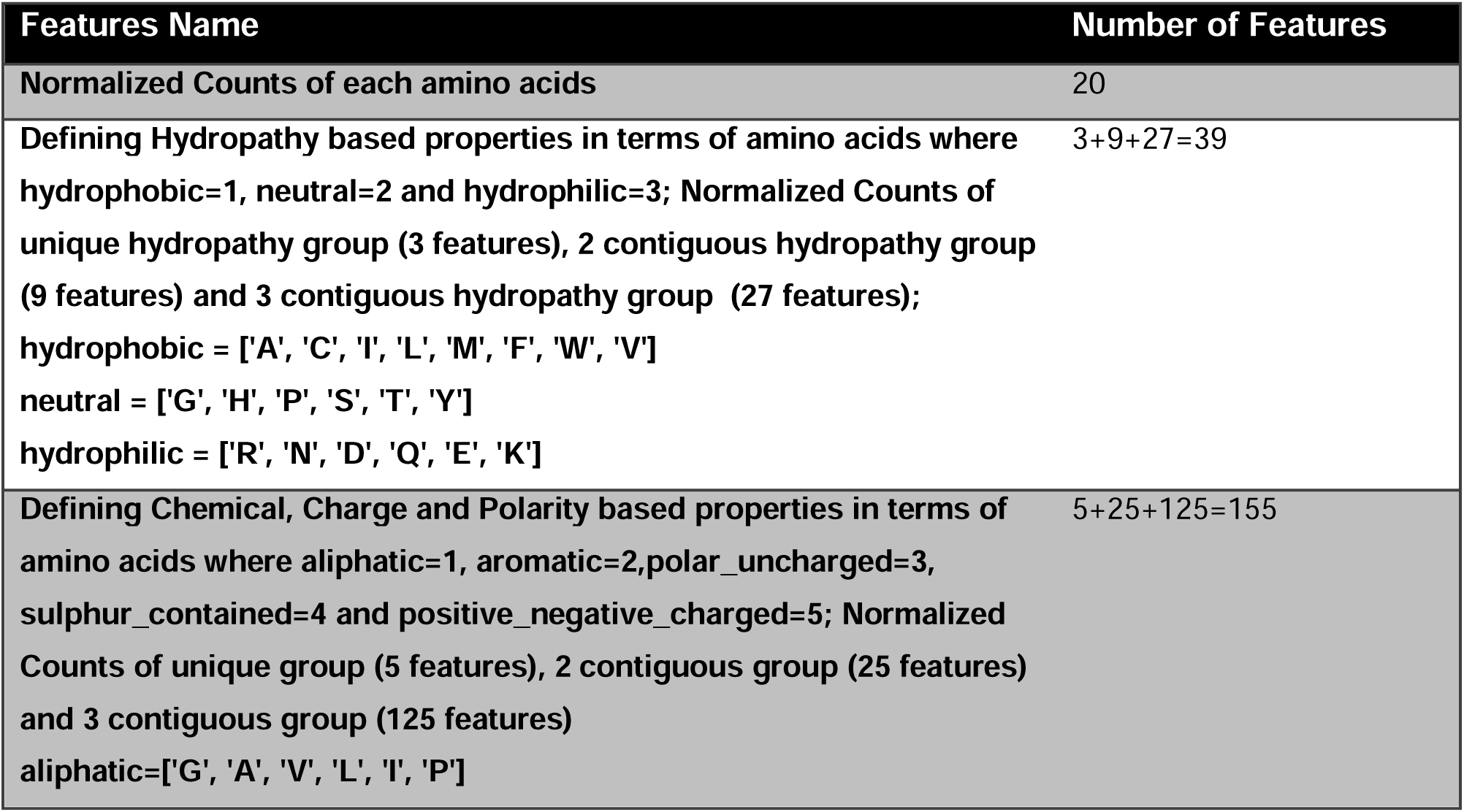
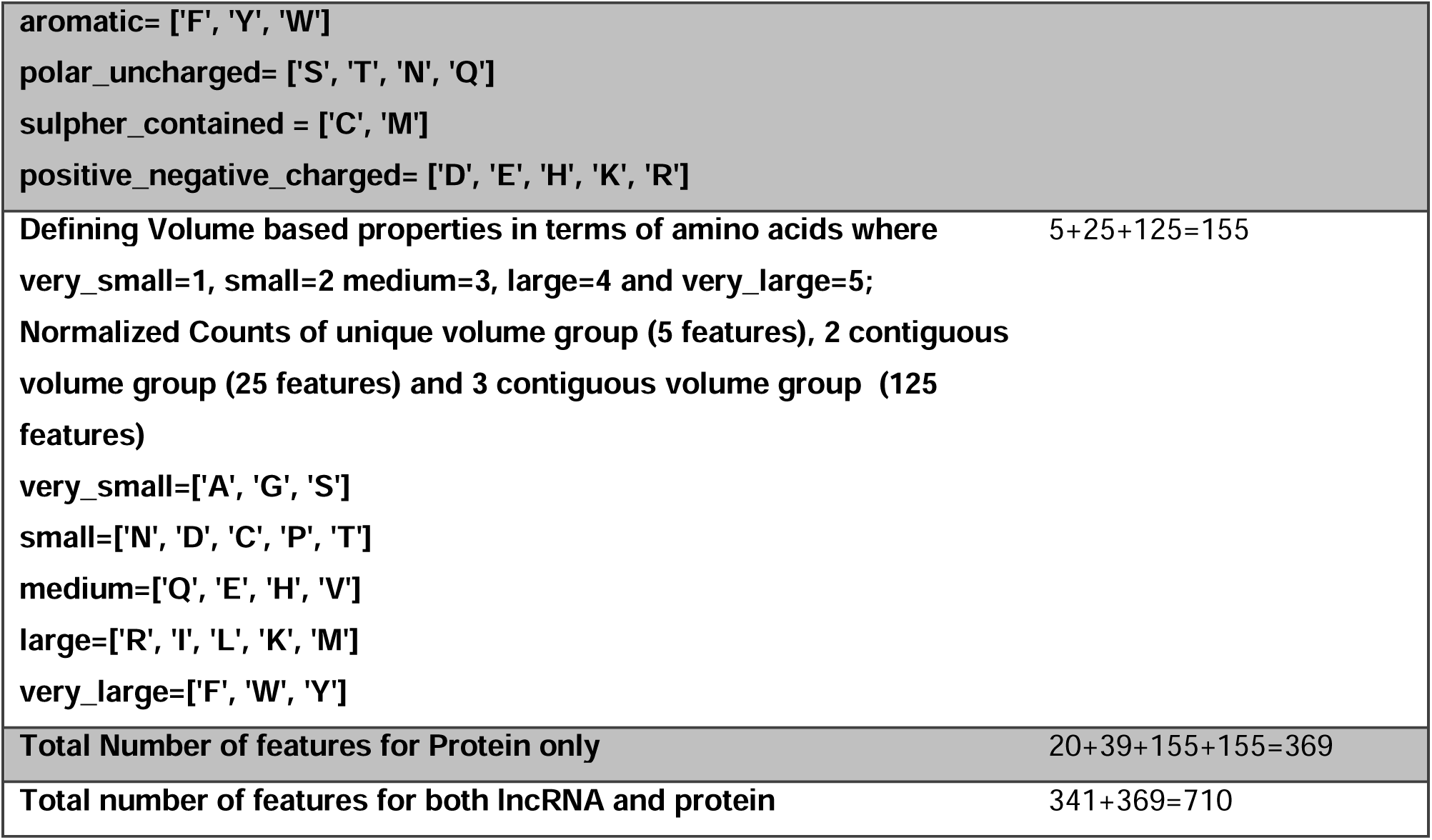
Detailed Feature Description extracted from raw protein sequences and cumulated number of lncRNA and protein features.

### Data Normalization

The set of features generated above, belong to varied ranges, which would prevent the ML models for smoother convergence. Normalization procedure would scale those diverse ranges such that they become comparable resulting in removal of unwanted bias corresponding to larger variables. Following equation (1) provides the formulation behind the normalization procedure.

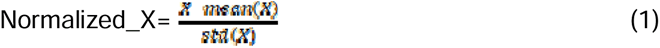

We have implemented scikit-learn based StandardScaler method for normalizing the data (http://scikit-learn.org/stable/modules/generated/sklearn.preprocessing.StandardScaler.html) as provided in equation(1).

### ML Model Creation & Validation

Figure 4 showing the detailed flowchart corresponding to our ML model building phase where we have incorporated **Stacking Ensemble (StackingClassifier)** based model building architecture. This is achieved by incorporating various ML models across multiple layers. It follows more generalized approach by comparing with bagging (33) and boosting (34) strategies. Our model architecture have been segregated into two layers: **Layer 1** constitutes hyperparameter optimized models **LightGBM** (**35**), **XGBoost** (**36**) & their variants whereas, **Layer 2** merges the output from previous layers base models using **Logistic Regression** based meta-learning technique(shown in Figure 4).

**Figure 4:**
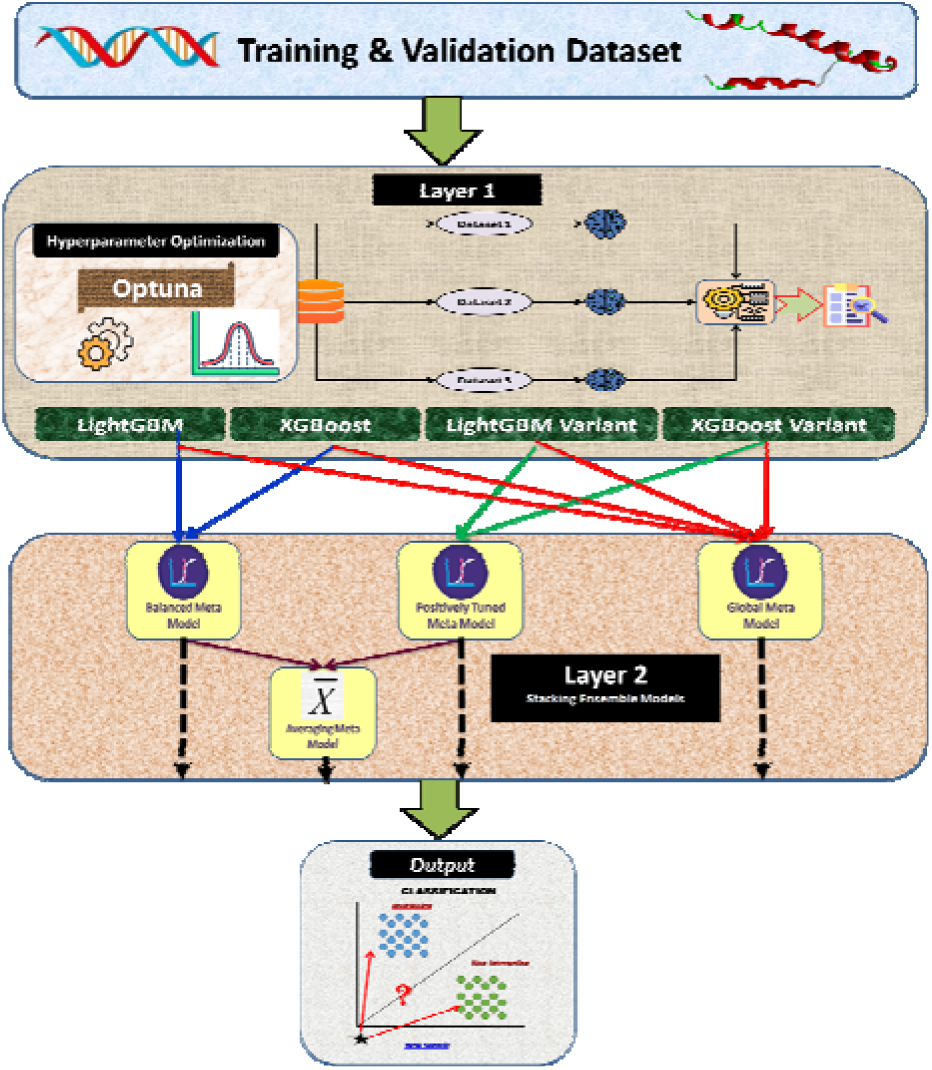
Machine learning model building. This figure elaborates the set of hierarchical layers associated with various machine learning models utilized in LncPTPred tool.

#### Layer 1 (Base Model)

Layer 1 constitutes of 4 base models LightGBM, XGBoost and their corresponding variants. It has been found that both the general LightGBM & XGBoost models possess biasness towards non-interactive output, thus we have tweaked the associated weights a little bit in order to reduce such kind of biasness, resulting in formation of model variations.

#### Layer 2 (Stacking Ensemble Model)

Layer 2 is also known as Stacking Ensemble layers, constructed by combining the prediction output extracted from the base model of previous layers and redirected to Logistic Regression which also acts our meta-learner to generate final prediction output. Here we have built four stacking meta-learner models by incorporating various combination of the base model as detailed in Figure 4**. Balanced Meta Classifier,** which merges the prediction output from general optimized LightGBM and XGBoost are fed into Logistic Regression meta-learner. **Positively Tuned Meta Classifier,** combine the outputs from variants of both LightGBM and XGBoost into the Logistic Regression meta-learner. **Averaging Meta Classifier** doesn’t create any separate model, rather takes prediction probabilistic output from **Balanced Meta Classifier** & **Positively Tuned Meta Classifier.** It performs averaging operation and gives separate output. Finally, **Global Meta Classifier** combine prediction outputs from all of the 4 base models from Layer 1 i.e. LightGBM, XGBoost & their corresponding variants and send them to the Logistic Regression meta-learner in Layer 2.

#### Hyper Parameter Optimization (HPO)

HPO depicts fine-tuning of various hyper-parameters associated with all kind of ML models. We have incorporated python based tool **Optuna** (**25**) which internally utilizes Bayesian Optimization (37) by employing efficient sampling and pruning algorithm to fine tune the hyperparameters from dynamically constructed search space. This tool has been incorporated upon **LightGBM**, **XGBoost** and their variant models to optimize their respective hyper-parameters for execution of lncRNA-protein target prediction task.

#### Performance Metrics

In order to validate the efficacy of LncPTPred tool against various external tools, multiple performance metrics have been taken into consideration consisting of **Accuracy, Sensitivity, Specificity, F1 score, ROC-AUC** & **Confusion Matrix.** LncPTPred performs binary classification task having discrete target variable containing 1 and 0 annotating positive and negative interaction respectively.

#### Comparison with other Validated Models

In order to validate the efficacy of LncPTPred tool, we have compared the performance of our model against 3 tools **RPISeq, LPI-CNNCP** & **Capsule-LPI.** Non-availability of pre-trained models (https://github.com/plhhnu/LPI-HyADBS; https://github.com/mrzResearchArena/PyFeat/), deprecation of the python environment (13) limited us to include others in the comparison. Further the web-pages of some tools (10,12) are unavailable.

Constructed using traditional ML models, RPISeq is one of the well cited tools defined for the prediction task of lncRNA-Protein interaction consisting of Support Vector Machine (SVM) and Random Forest (RF) where the features are defined using 3-mer protein and 4-mer RNA extracted from the sequences. Rest of the two tools have incorporated deep learning based prediction model. LPI-CNNCP proposed the Convolution Neural Network (CNN) model to perform prediction task where the feature set was generated using copy padding trick to convert variable length sequences of lncRNA/protein into fixed length inputs, further incorporating one-hot encoding in order to transform CNN-compatible image data. Capsule-LPI tool embeds multimodal features in terms of secondary structure, motif, sequence and physiochemical properties where the Capsule network forms the basis of their prediction model. From the result section it can be easily visualized that LncPTPred completely outperforms these specified models.

#### Webserver Implementation and generating the output

Both **Python** & **R** programming languages have been incorporated in order to execute the entire **LncPTPred,** lncRNA-Protein target prediction task. The software packages associated with Python language (Version 3.8) have been downloaded using **Anaconda** (https://www.anaconda.com/) suite whereas **R** based packages have been downloaded using **cran** (https://cran.r-project.org/) and **Bioconductor** (https://www.bioconductor.org/) (**30**) tools. Sequences corresponding to raw RNA fragments obtained from CLIP experiments are generated using **Bioconductor** based **BSgenome** (https://bioconductor.org/packages/BSgenome/) package. In order to perform the entire data wrangling operation corresponding to target prediction task we have utilized both **pandas** (https://pandas.pydata.org/) (Version 1.3.3) and **numpy** (https://numpy.org/)(Version 1.19.5) tools. As per as the machine learning models are concerned we have utilized **scikit-learn** (https://scikit-learn.org/) (Version 1.0.2) package to execute data normalization, formation of stacking ensemble model and performance based metrics generation. **LightGBM** (https://LightGBM.readthedocs.io/) (Version 2.3.1), **XGBoost** (https://xgboost.readthedocs.io/en/stable/python/index.html) (Version 1.6.2) have been deployed for creation of ML models whereas, **Optuna** (https://optuna.org/) performs the hyperparameter optimization associated with those models. Data visualization has been encompassed by both **Matplotlib** (https://matplotlib.org/) in python and **ggplot2** (https://ggplot2.tidyverse.org/) in R. **Logomaker** (**38**) (https://logomaker.readthedocs.io/en/latest/) (Version 0.8) is utilized for sequence visualization in terms of sequence logo. We have utilized **HTML5, Javascript** and **PHP** to build the web-server.

In order to perform prediction analysis user needs to select protein names from given list & provide the lncRNA sequence. Window length section specifies the fragmented length of given lncRNA sequence where min value is 10, max value is 30 and default is 20 as determined by violin plot in Figure 3 where length distribution of dataset mostly lies between 20 and 25. Shift size determines moving window length where default setting is considered to be 1, as our main aim is to include maximum overlapping window. Finally, the strand is to be considered as + / – .

Given the protein name, lncRNA sequence, window size, shift size and strand information provided by the user, the results are generated in following steps:

- LncRNA sequence is segmented according to the window size and shift size.
- Corresponding to each segment, if prediction probability≥0.5 predicted by at least 3 of the 4 models, it is considered as interacting segment else not.
- Finally, the overlapping segments are merged to get the **Final_Interacting_Score** which denotes the interaction probability corresponding to each of the binding segments of a particular lncRNA.

## RESULTS & DISCUSSIONS

At first we have checked for specific sequence motifs (contained within the nucleotide sequence corresponding to the RNA binding proteins) which affects lncRNA’s binding affinity. Thereafter we have performed a set of experiments to evaluate the performance of LncPTPred.

*Specificity of an LncRNA target binding site*: Previous reports suggest greater affinity of an lncRNA towards AU rich region for HuR (39), FUS (40) and AUF1(41). The positive interacting nucleotide sequences (corresponding to these RBPs) extracted from such RBP specific CLIP-Seq experiments were scanned and similar motifs were observed. Figure 5 showing the sequence logo represents the predominance of AU(T)-rich region at the binding location of the 3 RBPs with the lncRNA segments.

**Figure 5:**
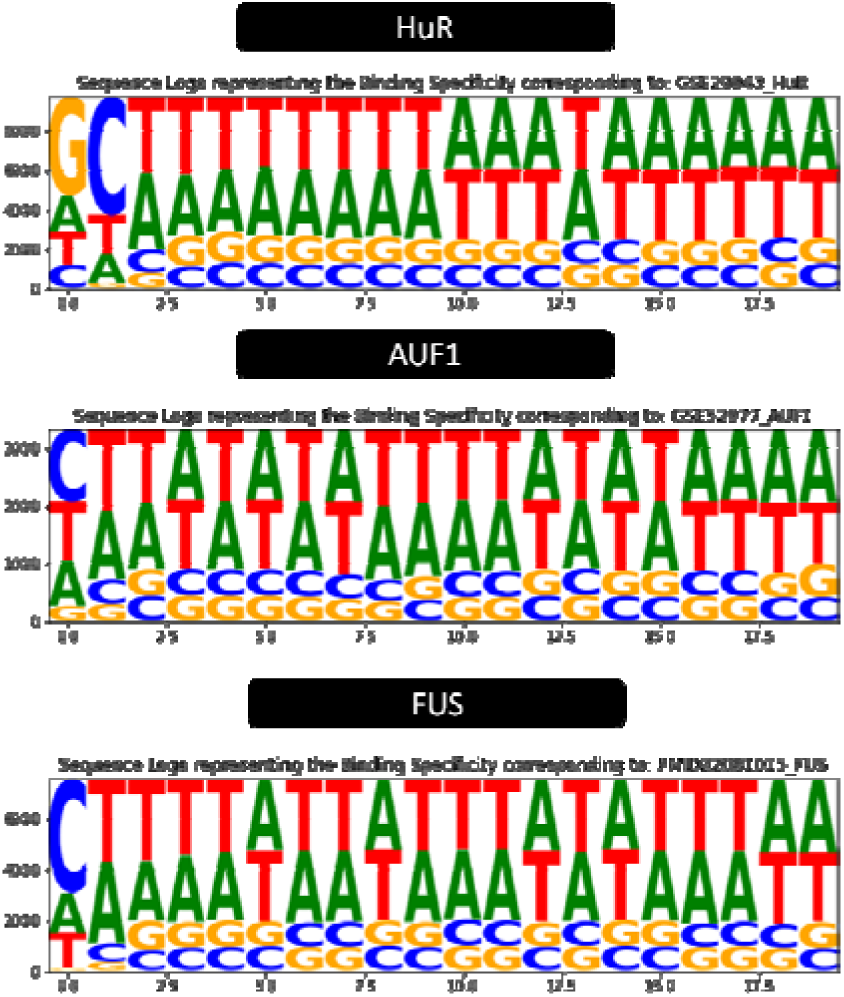
Sequence logo plot describing the binding specificity of lncRNAs with proteins. Logo plots showing the binding specificity corresponding to positive interacting lncRNA segments associated with 3 proteins HuR, AUF1 and FUS.

*Performance of LncPTPred for lncRNA-protein target prediction*: We have implemented stacking ensemble based ML models consisting of 4 separate classifiers **Balanced Meta Classifier, Positively Tuned Meta Classifier, Averaging Meta Classifier** & **Global Meta Classifier.** Various metrics have been utilized to measure the efficacy of LncPTPred tools, viz. **Accuracy, Sensitivity, Specificity, F1 score, ROC-AUC** & **Confusion Matrix.** In this paper we have compared the performance of the abovementioned 4 models used in **LncPTPred** with 2 models used in **RPISeq** viz. Support Vector Machine (SVM) and Random Forest (RF), **Capsule-LPI** & **LPI-CNNCP** tools using both validation and unknown test data. Figure 6a **& 6b** shows the bar diagram comparing the Accuracy, F1 score, Sensitivity and Specificity of LncPTPred models with RPISeq, LPI-CNNCP & Capsule-LPI models for validation data and unknown test data respectively.

**Figure 6:**
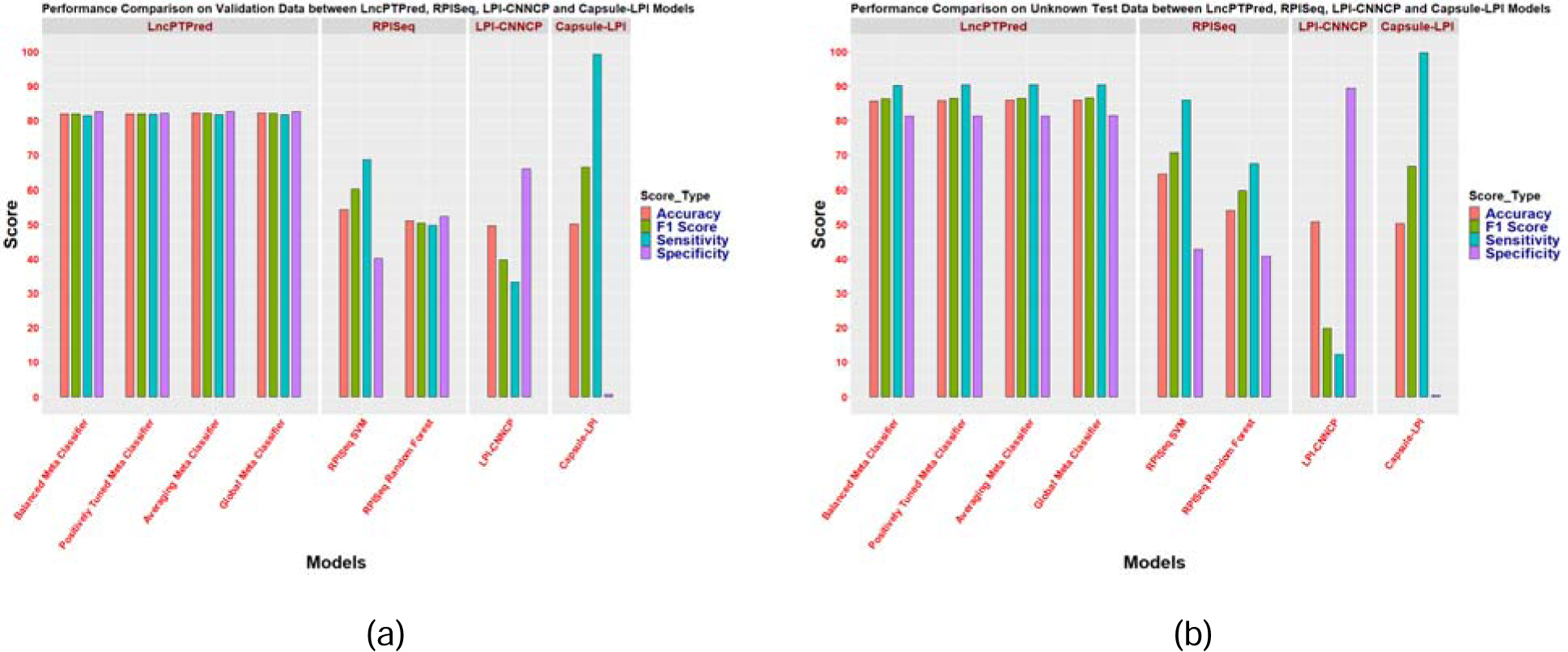
Performance Comparison of the various ML Models used to develop LncPTPred tool with RPISeq, LPI-CNNCP and Capsule -LPI for (a) Validation and (b) Unknown Test data. Bar chart showing the performance comparison among various LncPTPred models (Balanced, Positively Tuned, Averaging & Global Meta Classifier) with RPISeq models (SVM and RF), LPI-CNNCP and Capsule-LPI in terms of Accuracy, F1 Score, Sensitivity and Specificity metrics corresponding to (a) Validation and (b) Unknown Test data.

Accuracy and F1 score of the models used for LncPTPred outperforms the other models for both the validation and unknown test datasets. On the contrary, sensitivity score of Capsule-LPI is 100 whereas its specificity score is 0 (for both the datasets) which reveals the biasness of Capsule-LPI towards positive prediction leading towards greater false positive outputs. Further, the specificity score of LPI-CNNCP is very high and sensitivity very low (for both the datasets) which reveals the biasness of LPI-CNNCP towards negative prediction leading towards greater false negative outputs. This is better revealed in the confusion matrix for validation and unknown test data provided in Figure 7a **& 7b.** Hence, considering the global metrics like Accuracy and F1-score, LncPTPred scores the highest as compared to LPI-CNNCP, Capsule-LPI and RPISeq.

**Figure 7:**
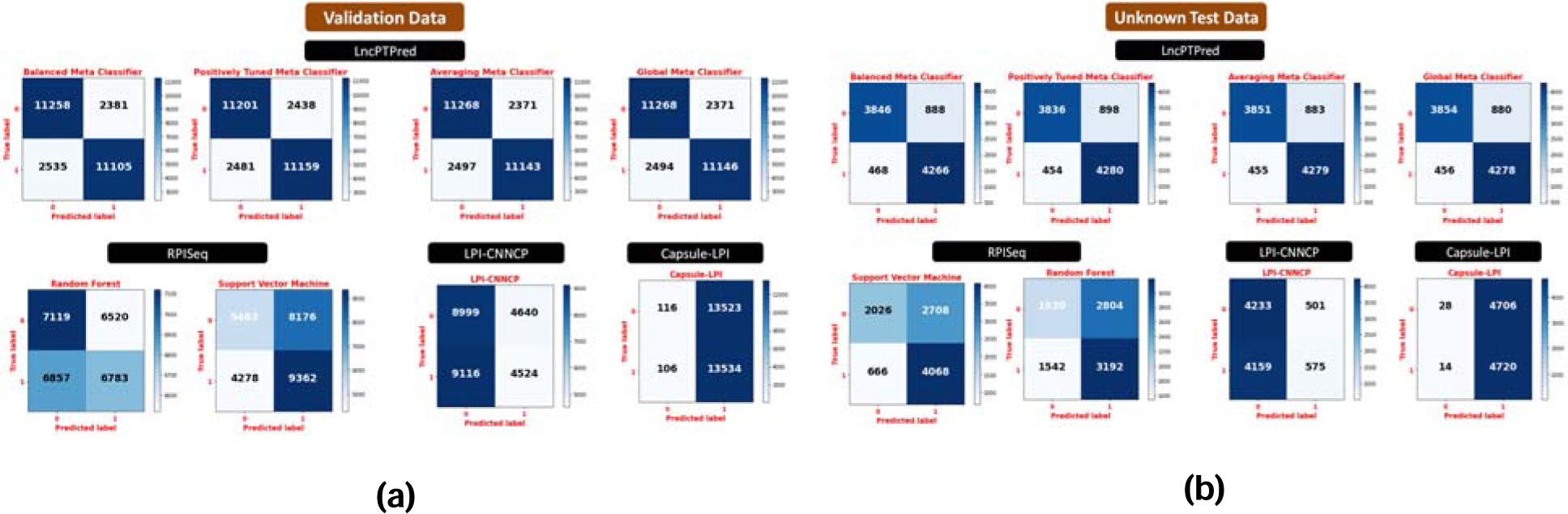
Confusion Matrix showing comparison of Performance Statistics corresponding to LncPTPred tool with RPISeq, LPI-CNNCP and Capsule-LPI for (a) Validation and (b) Unknown Test data. This Matrix shows the True Positive & True Negative predictions representing the number of accurate predictions whereas False Positive & False Negative predictions representing the number of wrong predictions comparing LncPTPred tool (viz. Balanced, Positively Tuned, Averaging & Global Meta Classifier) with RPISeq (viz. SVM and RF), LPI-CNNCP and Capsule-LPI tools.

Same holds true for ROC-AUC curve (as shown in Figure 8a **& 8b)**, where the models used in LncPTPred covers maximum area (>0.9) for both validation and unknown test data as compared to that covered by other classifiers.

**Figure 8:**
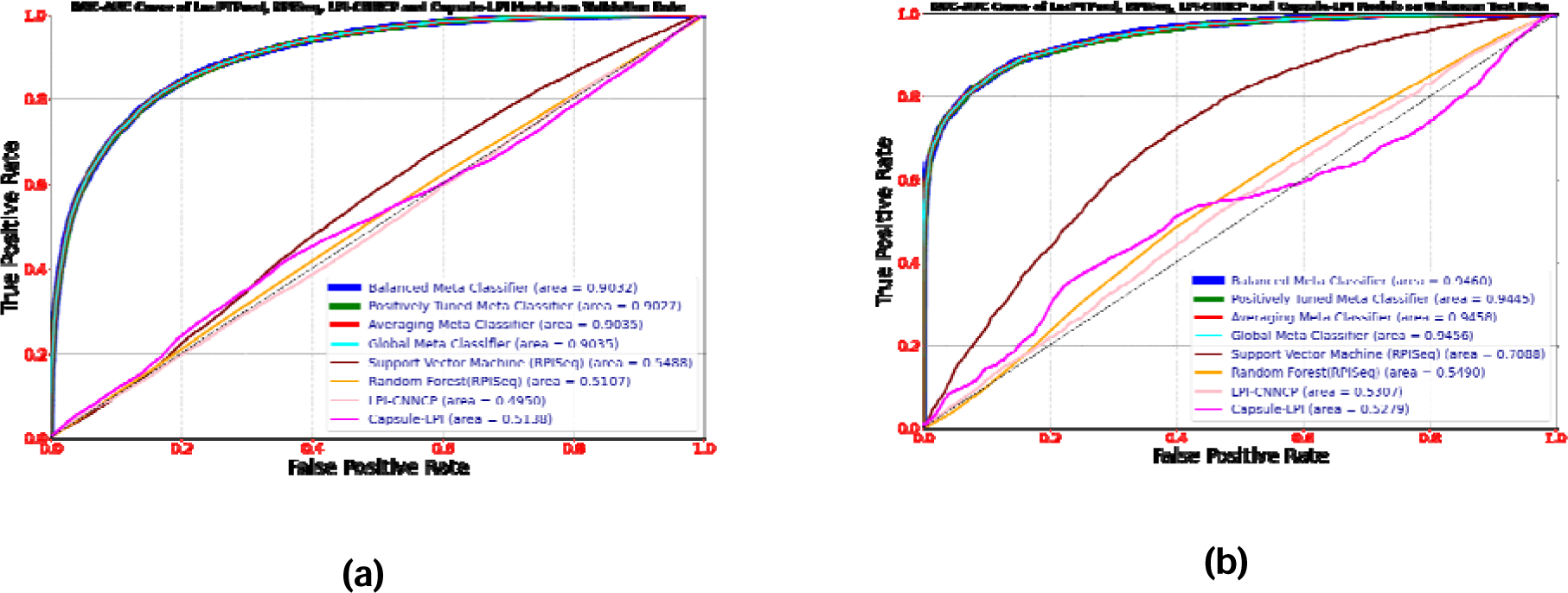
ROC-AUC Plot for (a) Validation and (b) Unknown Test data. ROC-AUC curve depicting the performance of the classification models used for developing LncPTPred, RPISeq, LPI-CNNCP and CAPSULE-LPI corresponding to (a) Validation and (b) Unknown Test data

### LncPTPred performance corresponding to disease specific lncRNA-protein interactions

We have incorporated LncPTPred tool in the context of disease specific lncRNA-protein target prediction. Here we have considered experimentally validated disease specific lncRNA-protein target interactions tabulated in **Table 6**.

**Table 6:**
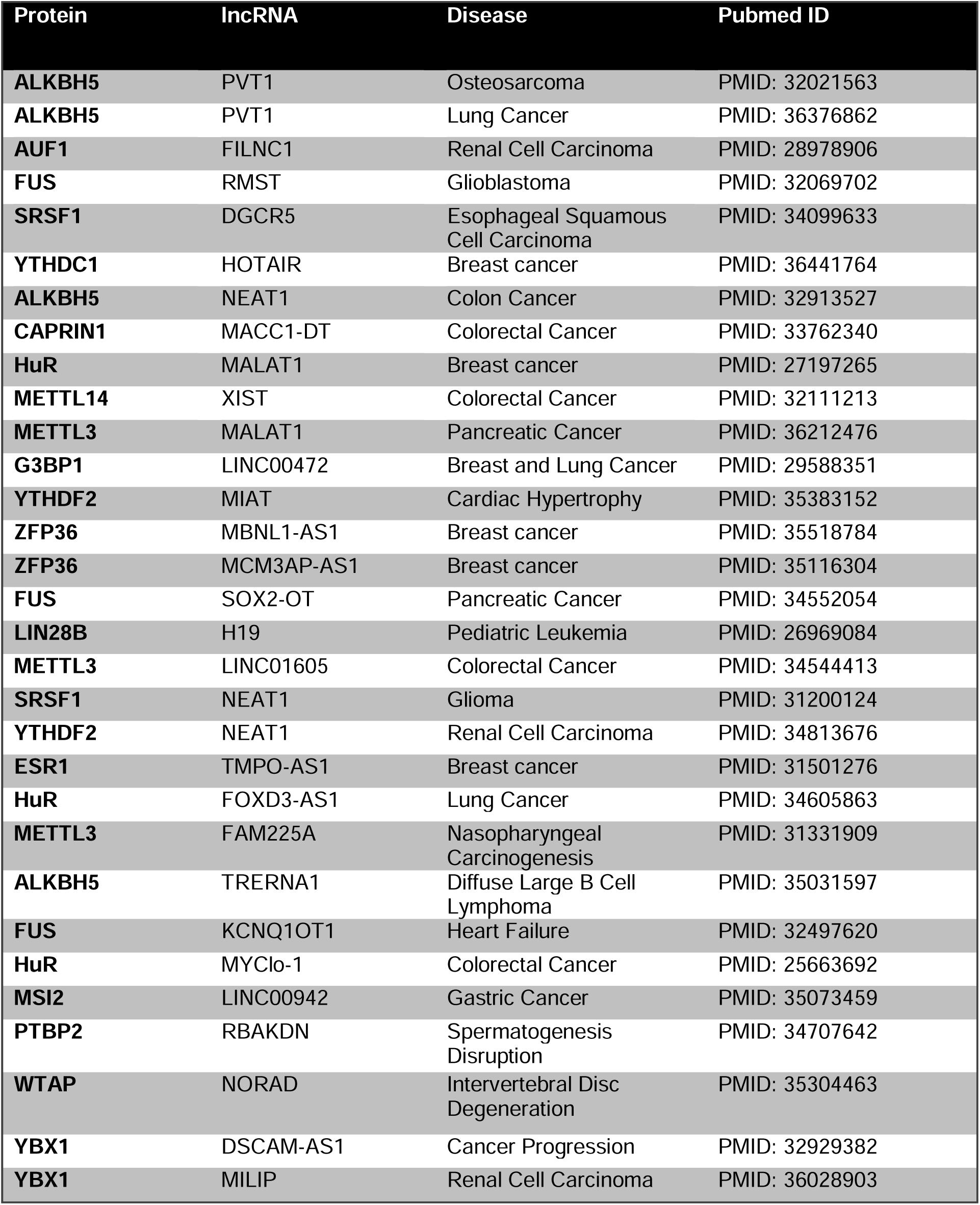
Disease specific reported lncRNA-Protein interaction.

Most of these lncRNAs have multiple transcripts. LncPTPred predicts lncRNA-protein interactions with all these transcripts. Thereafter, binding score for each of the transcripts for a particular lncRNA is obtained from the percentage of interacting segments among all the available segments (based on the window size = 20 and shift size = 1). The lncRNA transcript (for a particular lncRNA-protein interaction) with the highest binding score has been finally depicted in the bar graphs of Figure 9a-9d.

**Figure 9:**
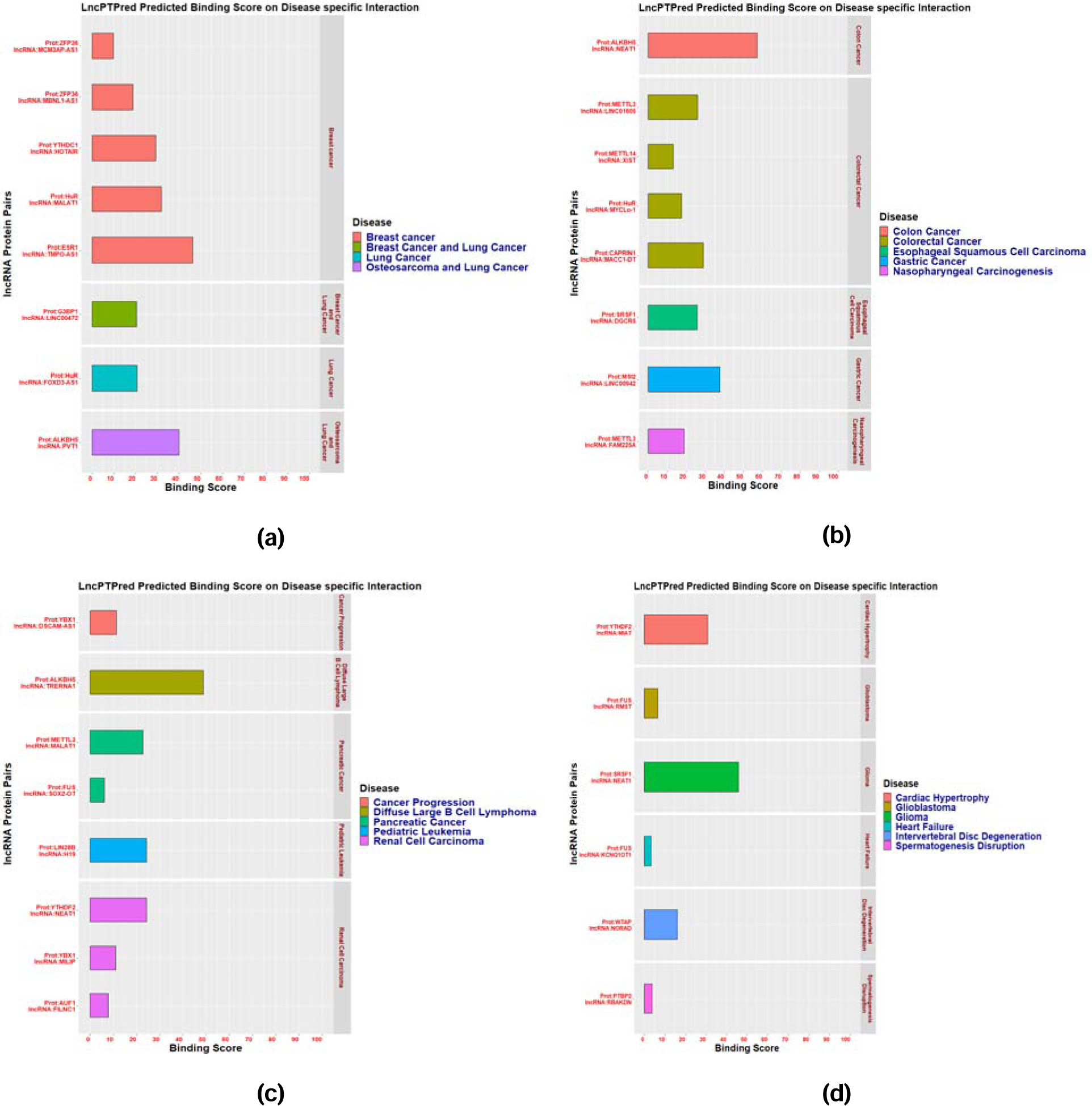
Performance of LncPTPred for predicting disease specific lncRNA-protein interactions. Horizontal Bar plot in figure (a)-(d) showing the lncRNA transcript with the highest binding score for a particular lncRNA-protein interaction validated in case of various diseases.

### Web Server Output

Figure 10a shows the user input page containing the input for lncRNA sequence, protein name and selection criteria regarding window size, shift size and strand specification. Once the user submits the query, it will provide the interacting segments and the **Final_Interacting_Score** as output. In order to show the binding specificity of the lncRNA segments, a sequence logo diagram is provided as shown in **Figure 10b**.

**Figure 10:**
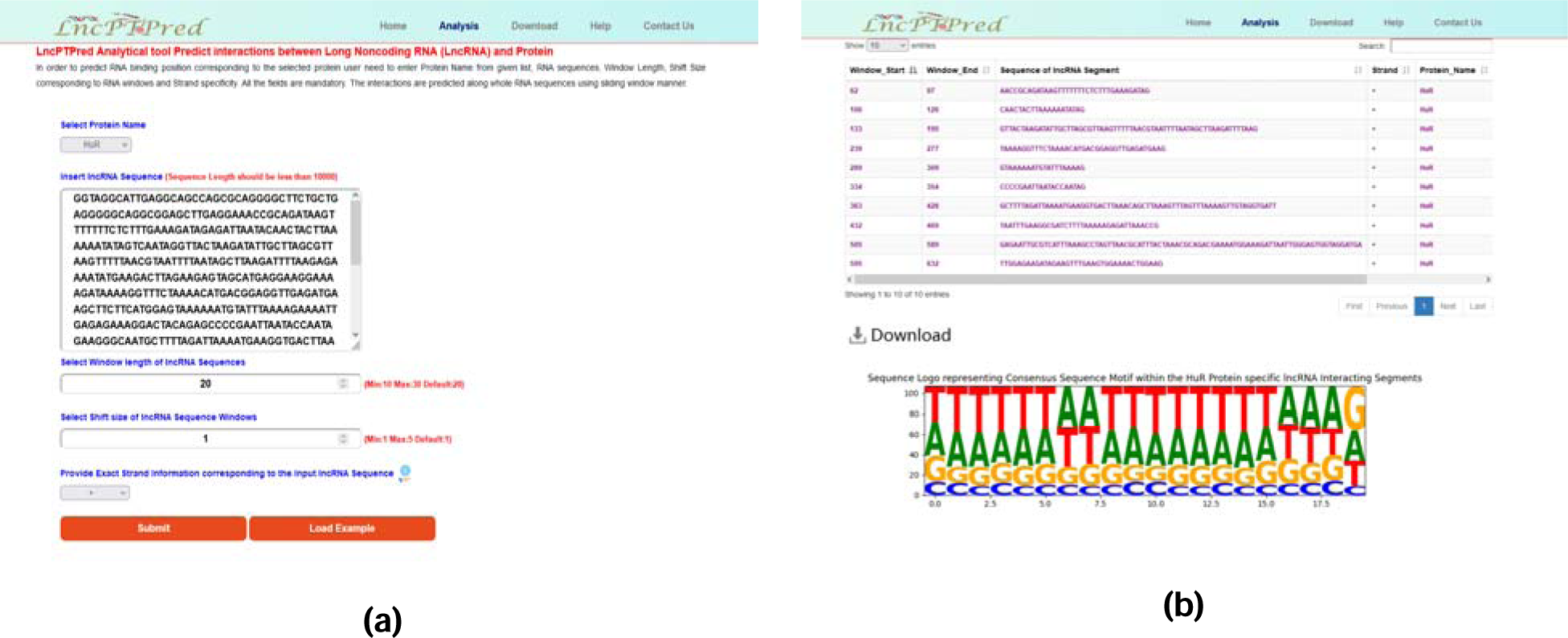
LncPTPred Web Server. Snapshot detailing the graphical user interface of LncPTPred web server (a) showing the required input data to execute the tool and (b) the output containing interacting lncRNA segment details, sequence logo diagram and the Final_Interacting_Score

## CONCLUSION

LncPTPred is an ML based target prediction tool involving lncRNA and protein. The biggest problem associated with this kind of target prediction scheme is the lack of real negative data. Recent reports on AI based lncRNA-protein target prediction algorithm relies on random shuffling techniques to tackle the issue ending up with noisy output resulting in most of the cases. We have tried to resolve the issue by considering CLIP-seq data. An extensive bioinformatics analysis have been done to extract sequence specific pattern associated with interacting positive data (corresponding to a particular protein) from within the CLIP-seq data. For that particular protein, the non-interacting segment served as the real negative data. We have considered Stacking Ensemble classifier as model architecture where Layer 1 consists of LightGBM, XGBoost & their variants as base model; the prediction output from the layer 1 have been merged using Logistic Regression meta-learner in Layer 2 to generate final output. Moreover we have enhanced the performance of our model by performing hyperparameter optimization using **Optuna** to set optimized values corresponding to model associated parameters. Finally, we have shown the efficacy of LncPTPred tool in two stages; comparing the performance of our models against RPISeq, LPI-CNNCP and Capsule-LPI tools where LncPTPred shows better performance against them. Moreover, we have showcased the efficiency of our model towards predicting some of the experimentally validated lncRNA-protein interactions across multiple diseases. Unlike the other state-of-the-art prediction tools, LncPTPred provides the exact location of interaction which will provide a big support for designing validation experiments. Further, on availability of more CLIP-seq datasets, we plan to upgrade this tool by providing predictions for more than 33 proteins in future.

## Supporting information

http://bicresources.jcbose.ac.in/zhumur/lncptpred/download_supple_file.php

## DATA AVAILABILITY

The detail information about raw CLIP-Seq data can be extracted from **Table 1** & **Supplementary File 1.** The data files associated with training, validation and unknown test data utilized for ML task can be downloaded from either the webserver http://bicresources.jcbose.ac.in/zhumur/lncptpred/download.php or from the actual github location https://github.com/zglabDIB/lncptpred/tree/master/dataset

## ACKNOWLEDGEMENTS

This work is funded by **National Network project of Bose Institute with Indian Statistical Institute and Vidyasagar University**, sanctioned by the Department of Biotechnology, Government of India, vide the sanction no. BT/PR40176/BTIS/137/84/2023 as well as Indian Council of Medical Research (ICMR) (sanction no. RBMH/FW/2020/10), Government of India.

## AUTHOR CONTRIBUTION

ZG hypothesized, conceptualized and provided overall supervision of the study. ZG and GD initially designed the experiments. Data curation and analysis was done by GD and TD. GD and ZG drafted the manuscript.

## SUPPLEMENTARY FILE LEGEND

**Supplementary File 1:** Detailed Information about PAR-CLIP & HITS-CLIP experiments considered for building the training, validation and unknown test dataset compatible for ML task.

